# Social structure defines spatial transmission of African swine fever in wild boar

**DOI:** 10.1101/2020.05.24.113332

**Authors:** Kim M. Pepin, Andrew Golnar, Tomasz Podgórski

## Abstract

1. Spatial spread of infectious disease is determined by spatial and social processes such as animal space use and family group structure. Yet, impacts of social processes on spatial spread remain poorly understood and estimates of spatial transmission kernels (STKs) often exclude social structure. Understanding impacts of social structure on STKs is important for obtaining robust inferences for policy decisions and optimizing response plans.
2. We fit spatially-explicit transmission models with different assumptions about contact structure to African swine fever virus (ASFv) surveillance data from Eastern Poland from 2014-2015 and evaluated how social structure affected inference of STKs and spatial spread.
3. The model with social structure provided better inference of spatial spread, predicted that ∼80% of transmission events occurred within family groups, and that transmission was weakly female-biased (other models predicted weakly male-biased transmission). In all models, most transmission events were within 1.5 km, with some rare events at longer distances. Effective reproductive numbers were between 1.1 and 2.5 (maximum values between 4 and 8).
4. Social structure can modify spatial transmission dynamics. Accounting for this additional contact heterogeneity in spatial transmission models could provide more robust inferences of STKs for policy decisions, identify best control targets, and improve transparency in model uncertainty.

## INTRODUCTION

Social [1] and spatial processes [2] are key drivers of pathogen transmission, yet their relative roles and influences on one another remain poorly understood [3, 4]. While both social structure and animal space use shape contact heterogeneity, they can have fundamentally different effects on contact structure and thus pathogen transmission dynamics [3-5]. For example, social organization into family groups (one dimension of social structure) causes local clustering of hosts that can alter pathogen transmission dynamics in a broad range of wildlife disease systems, such as malaria in primates, fungal infection in termites, intestinal parasites in African Artiodactylids, rabies in raccoons, bovine tuberculosis in badgers, and Chronic Wasting Disease in deer [6-12]. In contrast, animal space use mainly acts on overall connectivity in a population by limiting how far a disease can travel at each transmission event [13-16]. As such these processes are commonly modeled using different techniques and thus not accounted for in the same framework [3]. For example, social structure is often represented using network models [3, 17], while space use is often represented through distance functions describing how transmission probability changes with distance between hosts (spatial transmission kernels; STKs) [18]. STKs are key parameters for planning disease control strategies because they can be used to inform how surveillance and control strategies should be deployed spatially [2, 18-22]. However, without appropriately accounting for contact heterogeneities due to social structure, STKs could be uncertain or biased. Thus frameworks that can account for the effects of social structure on STKs are needed to better understand how different dimensions of contact heterogeneity shape pathogen transmission dynamics in wildlife populations and to optimize response plans using STKs.

Individual–based models (IBMs) are useful for incorporating social structure and space use separately in the same framework [7, 23, 24] to understand the significance of both processes on pathogen transmission dynamics. In general, IBMs can be especially useful for determining how much complexity is important to capture pathogen transmission dynamics well enough to effectively guide control policies [3, 25]. Previous work has used similar IBM structures to represent social structure and space use concurrently in systems as different as rabies in raccoons, African swine fever in wild boar, bovine tuberculosis in badgers, and foot-and mouth disease in feral swine to inform disease management strategies [7, 24, 26, 27], demonstrating how IBMs that are designed to account for individual-level variation in social and spatial parameters are widely applicable while allowing for an understanding of how individual-level nuances affect pathogen transmission dynamics. This flexibility is particularly well-suited for understanding how social structure modifies STKs. Secondly, social structure is usually temporally dynamic, which has been frequently neglected in data-driven social-network models due to the challenges with estimating changes in network structure over time. An IBM approach allows for natural incorporation of temporal changes in social network structure due to demographic changes.

To address gaps with understanding the impact of social structure on the spatial spread of disease, we developed an IBM fit to weekly surveillance data from wild boar in Poland from 2014-2015. In Eastern Europe, wild boar demonstrate limited spatial movement and cluster into family groups suggesting that both social structuring and space use are important for inferring the dynamics of African swine fever virus (ASFv) transmission [26, 28, 29]. Thus, the ASFv system provides an excellent opportunity to quantify effects of social structure on STKs and the magnitude to which model uncertainty could affect policy decisions using STKs. ASFv has been extremely difficult to control in wild boar partly due to a lack of robust predictions of spatial spread for risk-based mitigation [30]. In countries where ASFv is so widespread in wild boar such as Poland, robust estimates of STKs are crucial for improving response plans and risk assessment for domestic pig producers by enabling prediction of how fast and where ASFv will spread, and thus optimization of resource allocation across the landscape to surveillance and control. Also, understanding how social structure defines STKs can help to identify control targets that would minimize disease spread (e.g., the importance of controlling family groups versus adult males).

Without detailed genetic or contact tracing data to reconstruct transmission history, STKs are predominantly estimated indirectly by fitting pathogen transmission models to available case data [2, 18, 31]. Common assumptions of STK estimation methods include a single introduction event and perfectly observed data [32]. However, STK estimates could be biased if re-introduction events from outside sources spark epizootic foci in spatially distinct areas from the initial outbreak (i.e., being mistaken for transmission between clusters and causing artificially fat tails in the STK). Likewise, STKs could be biased if the surveillance system is biased towards detection of particular transmission events. Thus, methods that account for re-introduction and surveillance design are important in cases where ‘single introduction’ and ‘perfect detection’ assumptions are violated. Similar to many wildlife host-pathogen systems [31, 33-35], ASFv surveillance in wild boar is mostly passive [36] with only a small proportion of cases likely being detected (e.g., for ASFv in Poland: ≤ 30% of all ASFv carcasses detected by carcass surveillance, ≤ 1.7% of all active ASFv infections detected by hunter harvest, [26]), and genetic evidence suggests that at least two re-introductions from neighboring regions occurred during the time frame of our study [37].

In previous work we fitted models that accounted for the realities of multiple introduction events and partially observed data to ASFv surveillance data from wild boar in Poland to estimate the frequency of carcass-based transmission and re-introduction in outbreak dynamics [26]. This previous work predicted greater than 10 international introductions per year were necessary to sustain viral persistence at low host densities [26]. Here, we extended this modeling approach to understand how social structuring and space use assumptions affect estimates of STKs and spatial pathogen dynamics. We tested three different assumptions about space use and social structure: 1) neighborhood (local transmission only), 2) exponential decay (distance distribution that includes long-distance processes), and 3) distance distribution with social structure (social and spatial processes) (Fig. 1). We fit the different models to ASFv surveillance data using approximate Bayesian computation (ABC) and compared the model fits using distance metrics and R^2^. We then predicted the STKs under each set of assumptions using the fitted models and evaluated the effects of social and spatial processes on STKs and a key epidemiological parameter – the effective reproduction number. Our framework provides a general approach for understanding the relative role of social and spatial processes in pathogen transmission dynamics and quantifying STKs in the presence of realistic complexities (i.e., social structure, multiple re-introductions, partially observed case data).

**Fig. 1.**
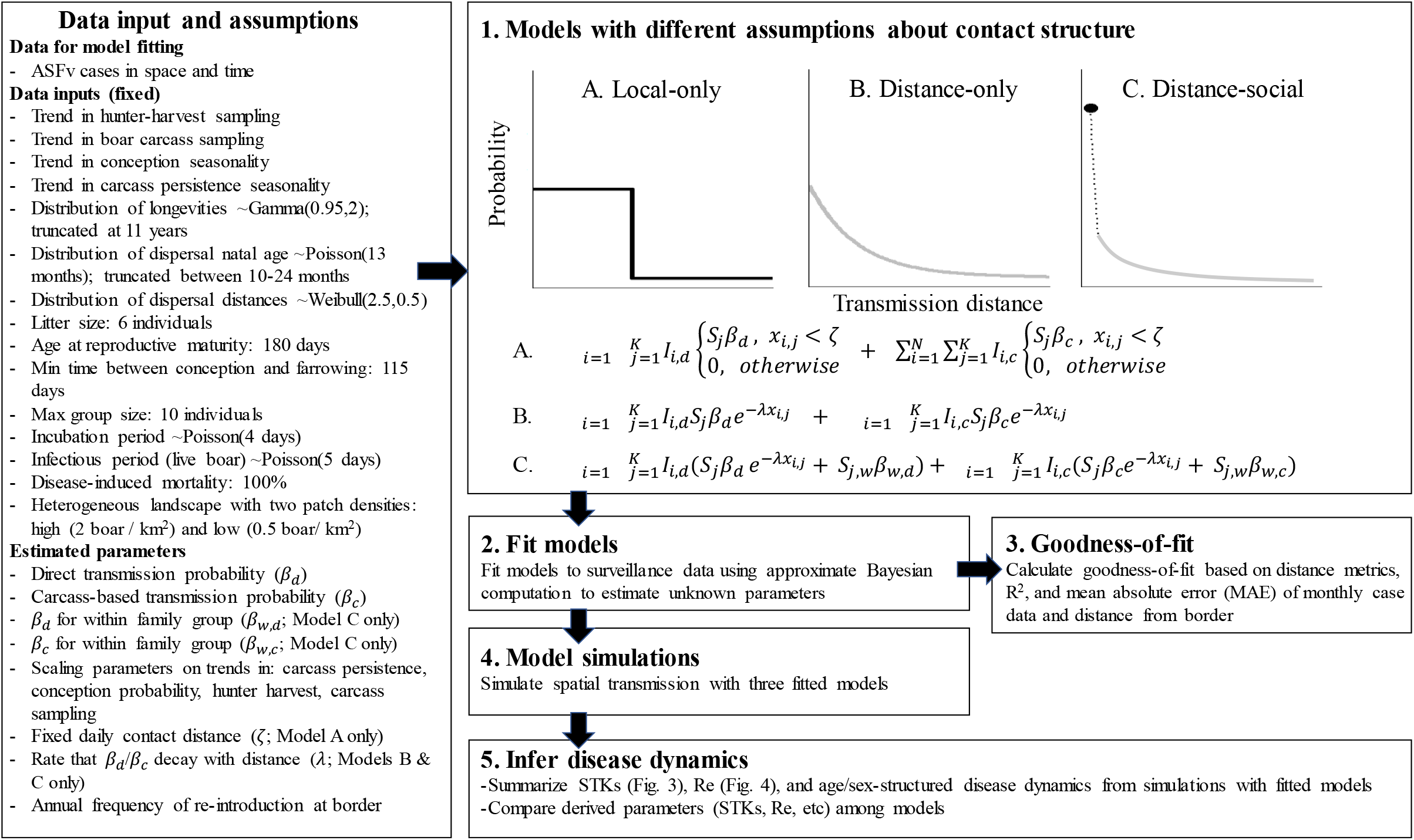
Schematic of methods and contact structures. Three different forms of contact structure were fitted to ASFv surveillance data (1. and 2.): A) neighborhood (local transmission only; local-only model), B) distance distribution (exponential decay that includes long-distance processes; distance-only model), and C) distance distribution with social structure (same as B. but also allows for different transmission probabilities for within versus between family groups; distance-social model). Goodness-of-fit was determined for each of the three fitted models (3.). Then the fitted models were used to simulate pathogen transmission dynamics (4.) and epidemiological metrics were derived from the simulated output (5.). *ζ* is a fixed local neighborhood (Eq. A) delimiting the contact radius, *x*_*i,j*_ is the distance between infectious individual *k* (*I*_*k*_) and susceptible individual *j (S*_*j*_*), α* is the rate at which transmission decays with distance (Eq. B and C), *d* denotes alive individuals or direct transmission, *c* denotes infectious carcasses or carcass-based transmission, β is the transmission rate that is specific to the transmission mechanism (*d* or *c*), and *w* denotes contact within the same family group whereas absence of *w* denotes contact among family groups (Eq. C).

## METHODS

### Study system and data

African swine fever virus (ASFv), a virulent virus of swine, emerged in domestic pigs in Georgia in 2007 following introduction from Africa [38]. After its initial emergence, the virus spread quickly to Eastern Europe becoming endemic in wild boar, which has challenged elimination. With no effective vaccine or treatment options, control strategies are focused on reducing swine movement, decontamination, and culling [39]. In wild boar, transmission occurs primarily through direct contact or contact with contaminated carcasses, consumption of contaminated food resources, and other forms of environmental or mechanical transmission are suspected [40]. Effectiveness of these strategies depends on being able to rapidly find new cases and target high-risk areas, thus models that can predict spatial spread are crucial tools. The index case of ASFv in wild boar was detected in February 2014 in north-eastern Poland (53°19’33”N, 23°45’31”E), less than 1 km from the border with Belarus. Subsequent cases occurred close to the Belarusian border [41, 42]. By the end of 2015, 139 wild boar tested positive for ASFv in the area, with maximum distance of 27.4km west of the border and a 100km range along the border. The affected area is dominated by a mosaic of woodlands and agricultural land (crop fields, pastures, meadows) with several large (several hundred square kilometers), continuous forests. On average, forest covers 53% of the area and wild boar densities range from 1.5 – 2.5 wild boar / km^2^ across all of Poland [43]. A total of 2470 samples from hunters and 205 samples from carcasses were collected from 8 administrative districts where ASFv occurred from Feb. 2014 through Dec. 2015. These samples were submitted to the National Reference Laboratory for ASFv at the National Veterinary Research Institute in Puławy, Poland for viral diagnostics. Surveillance data were used to fit the model and define spatio-temporal intensity of sampling for this analysis. We also used similar data from Jan. 2016 through Jul. 2016 to evaluate out-of-sample prediction. A detailed description of laboratory procedures and tests can be found in Woźniakowski et al. (2015) and Śmietanka et al. (2016).

### Process model

#### Overview

Previously we developed a spatially-explicit, individual-based model of ASFv transmission dynamics in wild boar that we fit to surveillance data using Approximate Bayesian Computation (ABC). This model accounted for: 1) social structure (independent adult males and matrilineal family groups that varied seasonally in size and composition due to birth rates, natal dispersal, and hunter harvest trends), and 2) spatial processes (daily space use and dispersal to new home range centroids), and was used to infer the role of carcass-based transmission in disease persistence. However, we did not investigate how social structure determines spatial transmission nor whether this level of complexity was necessary for capturing spatial transmission dynamics. Thus, here we used the same framework to evaluate three models that differed by social and spatial transmission process assumptions (described below) (Fig. 1). We estimated transmission parameters and some other epidemiological and demographic parameters as described below. With parameters from the fitted models we then predicted cases over time, spatial spread over time, STKs, effective reproductive numbers over time, and age- and sex-structure of infected individuals. All analyses were implemented in Matlab (Version R2016b, The MathWorks, Inc., Natick, Massachusetts, United States). A full description of the individual-based model and code for running it is given in Pepin et al. (2020). Below is an overview of the approach with emphasis on differences from our previous work.

#### Landscape

We used a gridded landscape to allow heterogeneity in population density across the landscape through density-dependent reproduction. We also allowed grid cell densities to affect dispersal (and thus potentially spatial spread of ASFv) by preventing dispersal to grid cells that were at carrying capacity. We chose a grid cell resolution of 5 x 5 km (25 km^2^) because it allowed fine-scaled heterogeneity in host density (close to home range size) while maintaining reasonable computation time. The total landscape size was 120 x 50 km (6000 km^2^), similar to the ‘infected’ zone in eastern Poland. Grid cells each had a carrying capacity of 0.5 or 2 boars/km^2^ (average density of 1.5 wild boar / km^2^), which is similar to previous estimates of local densities that were estimated to range from very low (<0.5 boar/km^2^) in poor habitats to high (>1.5 boar/km^2^) in high quality habitats [43]. This level of heterogeneous boar density fit the surveillance data better than homogenous densities of 1, 2 or 4 wild boar / km^2^ [26].

#### Attributes and demographic processes

Individual-boar attributes were monitored and updated at a daily time step. These included age, unique group identification, X and Y coordinates of the home range centroid, grid-cell ID; and status of life, reproduction, and infection. Thus, the distribution of wild boar locations was continuous but density was controlled at the grid-cell level. The variable attributes changed based on time, age, group size, grid-cell density, natal dispersal timing, and the pathogen transmission process. Fixed individual-level attributes included sex, dispersal distance, dispersal age, and age at natural death, which were all chosen from probability distributions [26].

Individual-boar status was updated by the following order of processes: daily movement (defined by the contact processes described below) and pathogen transmission, natural mortality (occurring according to the pre-set age), natal dispersal (occurring according to the pre-set age), dispersal due to other factors (i.e., family groups becoming too large, single females searching for groups; occurring based on current family group size), surveillance sampling (permanent removal of hunter-harvested individuals and carcasses), conception (rates dependent on current grid cell density), and new births (occurring with gestating females reach the end of their gestation period). Fixed parameters included longevity (a data-based distribution), litter size (6), age at reproductive maturity (180 days), minimum time between conception and farrowing (90 days), gestation time (115 days), age of natal dispersal (∼Poisson(13 months) truncated between 10-24 months), dispersal distance (∼Weibull(2.5,0.5)), maximum size of family groups (10), incubation period for ASFv (∼Poisson(4 days) truncated at 1), infectious period for ASFv (∼Poisson(5 days) truncated at 1), and disease-induced mortality (assumed to be fixed at 100% lethality for infectious individuals). There were also fixed seasonal trends that varied monthly for conception probability and carcass persistence that were based on data [44-47]. For example, seasonal birth pulses influenced group size and dispersal events over time (because dispersal was age-dependent), but we assumed that average daily contact distances were constant throughout the year. Rationale and sources for the processes and parameters were derived from ecological studies of wild boar and are described in Pepin et al. (2020).

#### Epidemiological states and processes

Epidemiological states for individual boar included: susceptible, exposed, infectious, infectious carcass, non-infectious carcass, and removed from the landscape. Mortality only occurred from the disease (leading to an infectious carcass) or reaching the age of longevity (leading to an non-infectious carcass). We assumed the disease was lethal in 100% of infectious individuals. The hunting process of alive individuals caused direct removal from the landscape (no carcass). Our model also included multiple spatio-temporal scales of spatial processes because the dispersal process (∼Weibull(2.5,0.5) allowed for longer-distance movements and occurred less frequently relative to the contact process that occurred daily and mostly at shorter distances.

We compared three different forms of contact structure: 1) neighborhood (local transmission only; local-only model), 2) distance distribution (exponential decay that includes long-distance processes; distance-only model), and 3) distance distribution with social structure (same as 2) but also allows for different transmission probabilities for within versus between family groups; distance-social model) (Fig. 1). We tested these three models because they represent increasing levels of ecological complexity in constraining spatial transmission. We viewed the local-only model as the coarsest representation of constraining transmission both spatially and within family groups. The local-only and distance-only models are common ways of considering contact in space at population-level scales [48], whereas the distance-social model incorporates heterogeneity due to social groups. For the local-only model, infectious individuals could transmit to all susceptible individuals within a fixed radius with equal probability. The radius of the local neighborhood was constant across individuals and time – thus similar to a queen’s neighbor effect (Eq. 1). For the distance-only model, infectious individuals could transmit to all susceptible individuals on the landscape, but the probability of transmission decayed with distance (Eq. 2). The distance-social model was the same as the distance-only model, except that transmission rates varied due to both group membership and space - individuals in the same family group had higher transmission rates with each other relative to those among family groups (Eq. 3). In general, the daily force of infection (*λ*) for each contact structure was defined as follows:

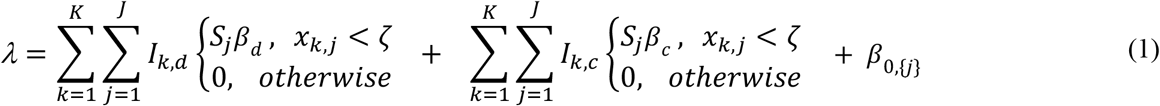

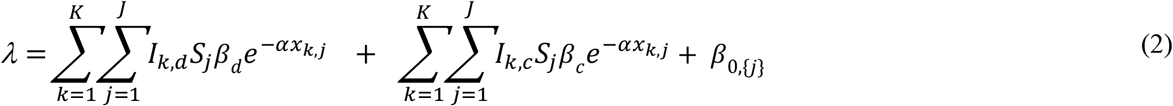

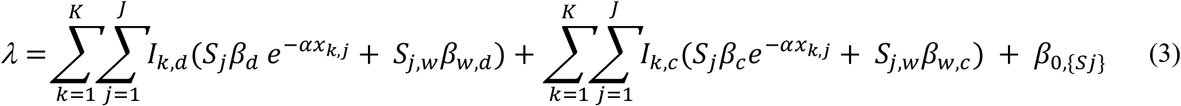

Where *ζ* is a fixed local neighborhood (Eq. 1) delimiting the contact radius, *x*_*i,j*_ is the distance between infectious individual *k* (*I*_*k*_) and susceptible individual *j (S*_*j*_*), α* is the rate at which transmission decays with distance (Eq. 2 and 3), *d* denotes alive individuals or direct transmission, *c* denotes infectious carcasses or carcass-based transmission, β is the transmission rate that is specific to the transmission mechanism (*d* or *c*), *β*_0,{j}_ is the baseline rate at which re-introduction occurs to susceptible individuals near the Eastern border ([49]), and *w* denotes contact within the same family group whereas absence of *w* denotes contact among family groups (Eq. 3).

### Observation model

Because surveillance sampling only tests a small proportion of the total population of wild boar in the region (< 2% monthly) it was important to calibrate the process model with an observation model. Thus, we sampled the true pathogen transmission dynamics according to the surveillance process that was used in Poland, i.e., alive individuals were available to be harvested by hunters, and carcasses were available to be found for carcass sampling; then both types of samples were evaluated to identify whether individuals/carcasses were infectious. As observed negative samples could not be georeferenced to the grid cell level (they were only available at the district level), we were not able to account for the spatial distribution of sampling accurately. However, we used the temporal trends in the surveillance data to determine the number of samples to collect per day by sampling the landscape at random but excluding wild boar < 6 months of age in hunter harvest (because they are typically not targeted by hunters; [50]) and wild boar < 3 months of age in carcass sampling (because they are unlikely to be found due to their small size and more rapid decay rates [51]). To determine the temporal sampling trends, we calculated the relative number of boar sampled by hunters and carcass-sampling from the data (number sampled on day *t*/maximum ever sampled separately for each method) to produce daily trends in the proportion of the population sampled. Then we multiplied the trend data for each method by the scaling factors (ρ_h_ and ρ_c_) to determine the daily proportion of boar that would be sampled (detection probability) by hunter harvesting or dead carcasses across the landscape at random.

### Model fitting and evaluation

Unknown parameters were estimated based on approximate Bayesian computation (ABC) with rejection sampling as described in Pepin et al. (2020). For all models, estimated parameters included: frequency of introduction at the eastern border (*β*_0,{j}_), *β*_*d*_, *β*_*c*_, scaling parameters on seasonal trends of hunted hosts (*ρ*_*h*_) and carcass sampling (*ρ*_*c*_), a scaling parameter on seasonal trends in the length of carcass persistence on the landscape (*π*), and a scaling parameter on seasonal patterns of host birth probabilities (*θ*). In addition, we estimated spatial parameters that describe three different contact structures: 1) ξ (nearest-neighbor), 2) *α* (the decay of contact probability with distance, and 3) *β*_*w,d*_ and *β*_*w,c*_ (direct and carcass-based transmission rates for within-group contacts). Prior distributions are listed in Table S1 (with restrictions: *β*_*d*_*>β*_*c*_, *β*_*w,d*_*>β*_*d*_, *β*_*w,c*_*>β*_*c*_) and were informed by movement and contact data [28, 29, 52, 53].

To sample across parameter space efficiently we used a Latin hypercube algorithm to generate 979,592 parameter sets and then ran the model twice on each parameter set (for a total of 1,959,184 iterations; or 2 chains of 979,592). *β*_*d*_, *β*_*c*_, and *ρ*_*c*_ were sampled on a log scale. To improve computational efficiency, a two-tiered approach was used to estimate posterior distributions of parameters. Simulations were terminated early if they were highly unrealistic compared to observed data, specific criteria and rationale were: 1) landscape-wide host density < 20% of the initial density because observed changes in wild boar density were only minor in the study during 2014-2016, 2) > 150 new cases per day because the maximum number observed per month was < 20; 3) no new cases sampled for 6 months because there was only 1 month with no cases detected after the first detection was made, or 4) > 300 total cases because that is more than double the actual number of observed cases. We then only considered parameter sets for which the simulation reached the end of the two-year time frame as candidate values for the posterior distributions. The posterior distributions consisted of all unique parameter sets (considering both chains) that were within the absolute distance of three metrics: the sum of absolute differences between observed and simulated surveillance data for monthly cases from live and dead animals (considered separately), and the maximum monthly Euclidian distance of cases from the eastern border. Distance metric tolerance values were 48 for monthly cases from carcasses, 24 for monthly cases from hunter-harvest samples, and 120 for maximum distance from the border. This allowed average error rates of 2 (carcass) and 1 (hunter harvest) cases, and 5 km from the border per month on average. These error rates represent levels of uncertainty that we expected from the data sources in our system, sensitivity analyses revealed that less stringent error rates would affect the posterior distribution estimates (data not shown), and more stringent error rates would require restrictively large computational resources unless prior distributions are more informed.

Average distance metrics for parameter sets from the posterior distribution were used to evaluate goodness of fit along with R^2^ values (squared correlation of observed and predicted case and spatial distance trajectories, Table S1) and mean absolute error (MAE, Fig. 2). The combination of these metrics were used for model selection and performance relative to one another, although we were also interested in comparing how the different model structures impacted parameter inference thus we examined output from all three models. For each fitted model we predicted outbreak dynamics using 100 random samples from the posterior distribution. The average of the 100 predictions was used to calculate R^2^ and MAE. We also tested the ability of our models to forecast ASFv dynamics by using the parameters estimated from fits to the 2014-2015 data to predict the first 7 months of 2016 (Jan.-Jul.). We predicted underlying STKs, effective reproductive number over time (R_e_), and age-sex structure of cases by simulating from the fitted models. Thus, STKs and R_e_ were model outputs. We calculated STKs through tracking the distance between each transmission event and summarizing the resulting distributions descriptively. Similarly, we calculated R_e_ by tracking the number of transmissions to new susceptible hosts that each live infectious individual and infectious carcass transmitted throughout their infectious periods and summarized daily R_e_ as means for individuals at the start of their infectious period.

**Fig. 2.**
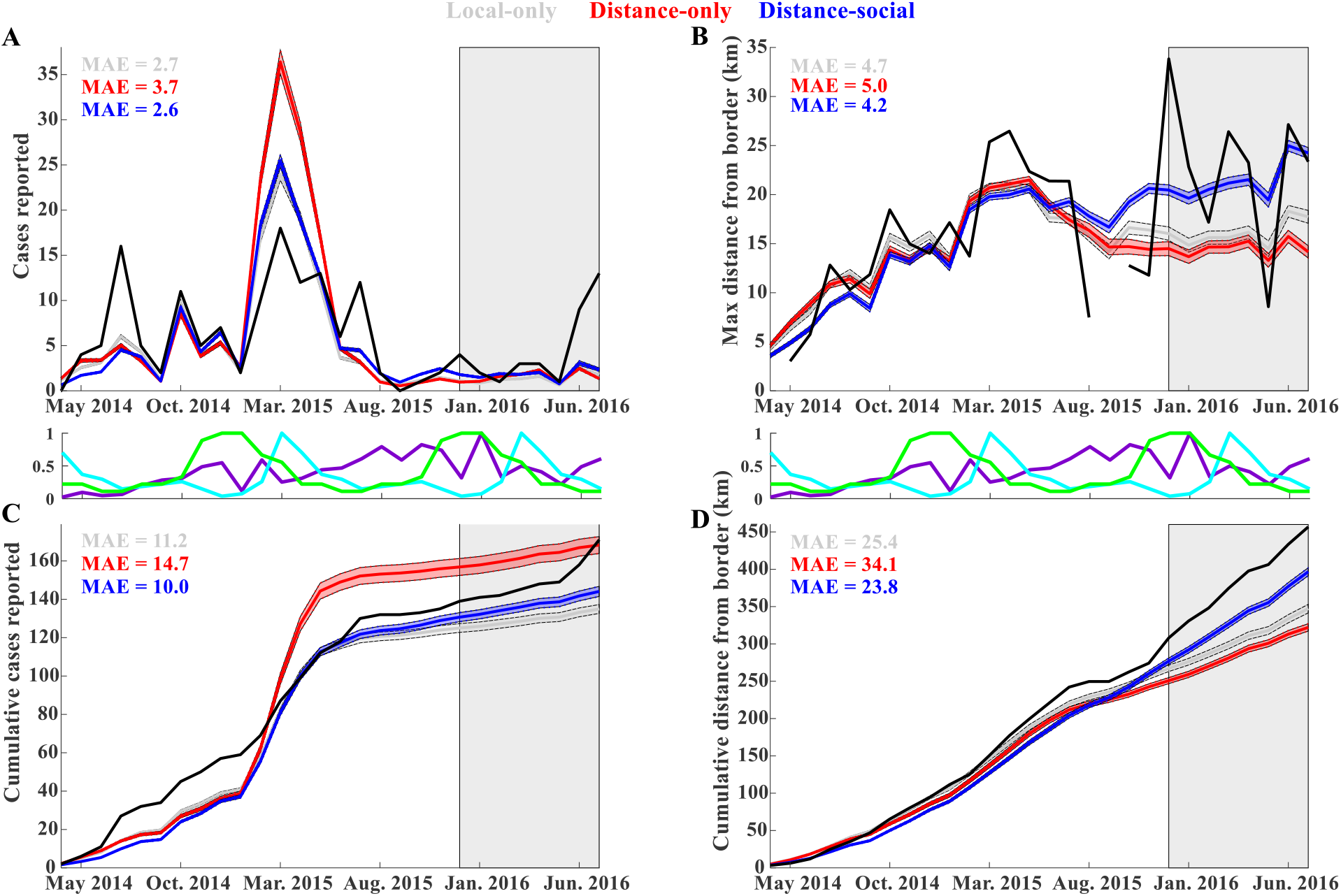
Model fits to the observed surveillance data (black). Lines are the mean predictions from 100 simulations with each fitted model (see legend in B for color code). Shading around the lines areas are 95% prediction intervals of the means. Shading on the right side of plots indicates the time frame for out-of-sample predictions. MAE is mean absolute error between the trajectories of observed and predicted monthly cases. Plots between the top and bottom panels show trends as a proportion of maximum values for total samples collected (purple), births (light blue), and carcass persistence time (green).

## RESULTS

### Parameter inference and model fit

The model with both social and spatial processes (distance-social model; Eq. 3) qualitatively captured spatial spread better than the local-only (Eq. 1) and distance-only (Eq. 2) models of spatial transmission, and the local-only model largely overestimated cases during the largest peak (Fig. 2). Also, the posterior distribution of transmission probabilities were much lower and more realistic for both distance models relative to the local-only model (see *β*_*d*_ and *β*_*c*_ in Table S1). The inferred STK for each model revealed two distinct peaks symbolic of within- and between-group transmission (Fig. 3) but predicted different amounts of within-group transmission when within- and between-group transmission probabilities were allowed to vary. The distance-social model predicted the highest amount of within group transmission (80%), followed by the distance-only (60%) and local-only (30%) models (Fig. 3). For both distance models, between-group transmission peaked between 0.5 km and 1 km, with a peak amount of transmission events reaching 5% and 2% for distance-only and distance-social models, respectively (Fig. 3). For the local-only model, between-group transmission events plateaued between 1-1.5 km at a frequency of 0.2 before dropping rapidly to 0 around 1.5km (Fig. 3). The inferred STKs for both distance models had long tails that indicated a low frequency of long-distance pathogen dispersal (Fig. 3).

**Fig. 3.**
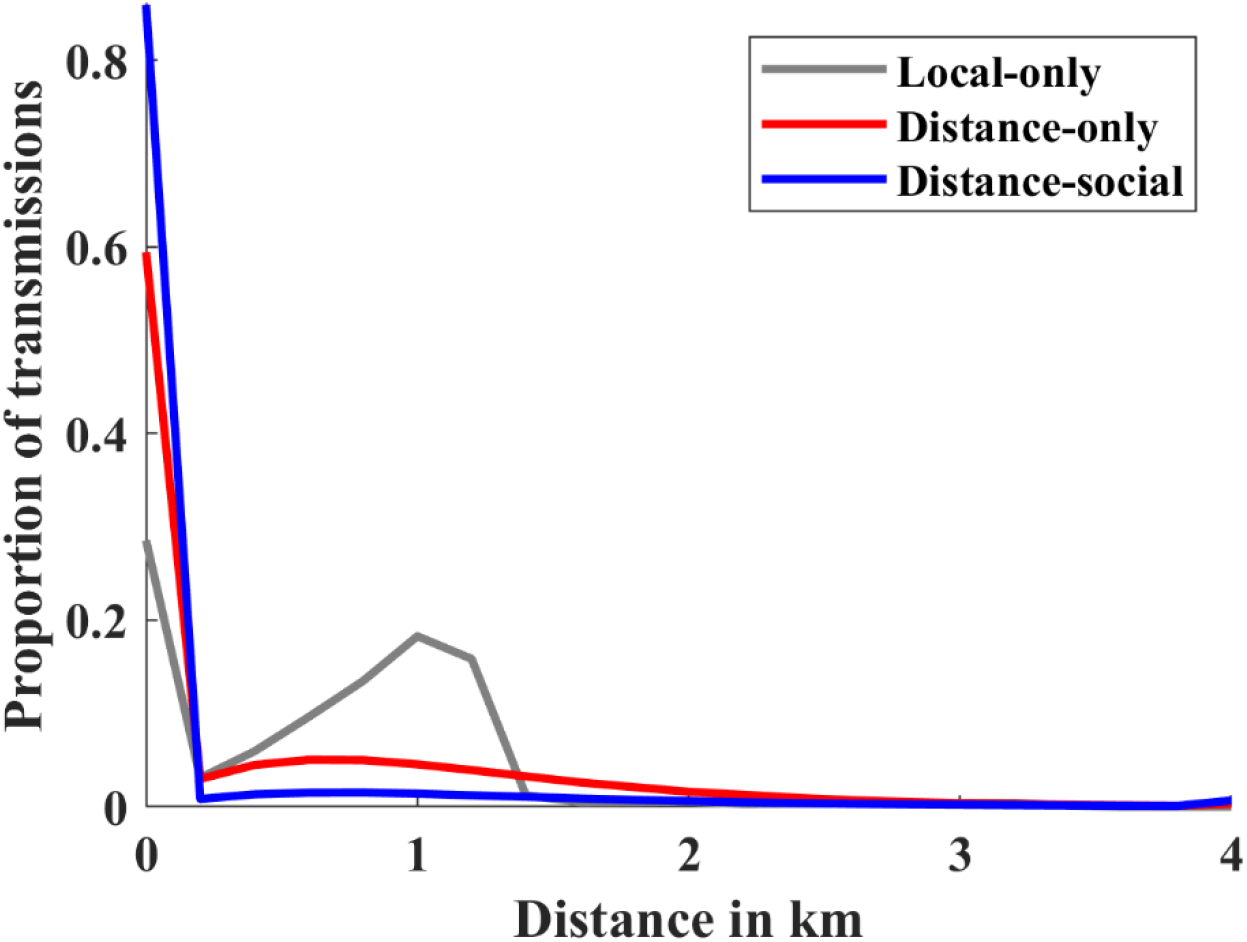
Inferred spatial transmission kernels (STKs) for each model (see legend). The X-axis is the distance between home range centroids of infectious and susceptible individuals for which transmission occurred. Y-axis is the proportion of all transmission events. Lines are the means of 100 simulations using random samples from the posterior distributions of the fitted models.

### Impacts of model structure on epidemiological processes

The specification of spatial and social transmission processes in the model structure resulted in different inferences of R_e_. The local-only model predicted higher average R_e_ over time (mean: 2.5 with 95% confidence interval: [1.8, 3.2]), followed by the distance-social (1.5 [1.1-2.0]), and then the distance-only (1.1 [1.0-1.3]) models, including both direct and carcass-based transmission (Fig. 4). However, predictions from the distance-only model suggested R_e_ is relatively homogenous over time, while the local-only and distance-social models predicted much more variability, with R_e_ values reaching above a value of 4 on multiple occasions, and above a value of 8 at least once (Fig. 4). The distance-social model predicted higher R_e_ during annual birth pulses (Fig. 4). The local-only and distance-only models predicted lower contributions of carcass-based transmission in overall R_e_ whereas the distance-social model predicted more similar levels of each transmission mechanism (with carcass-based transmission being slightly lower on average). All models predicted that the infected class is predominantly composed of juveniles (<6 months of age; Fig. S1-S3), reflecting the age-structure in the population. However, the local-only and distance-only models predicted a slight male-bias in infected individuals while the distance-social model predicted a slight female bias (Fig. S1-S3).

**Fig. 4.**
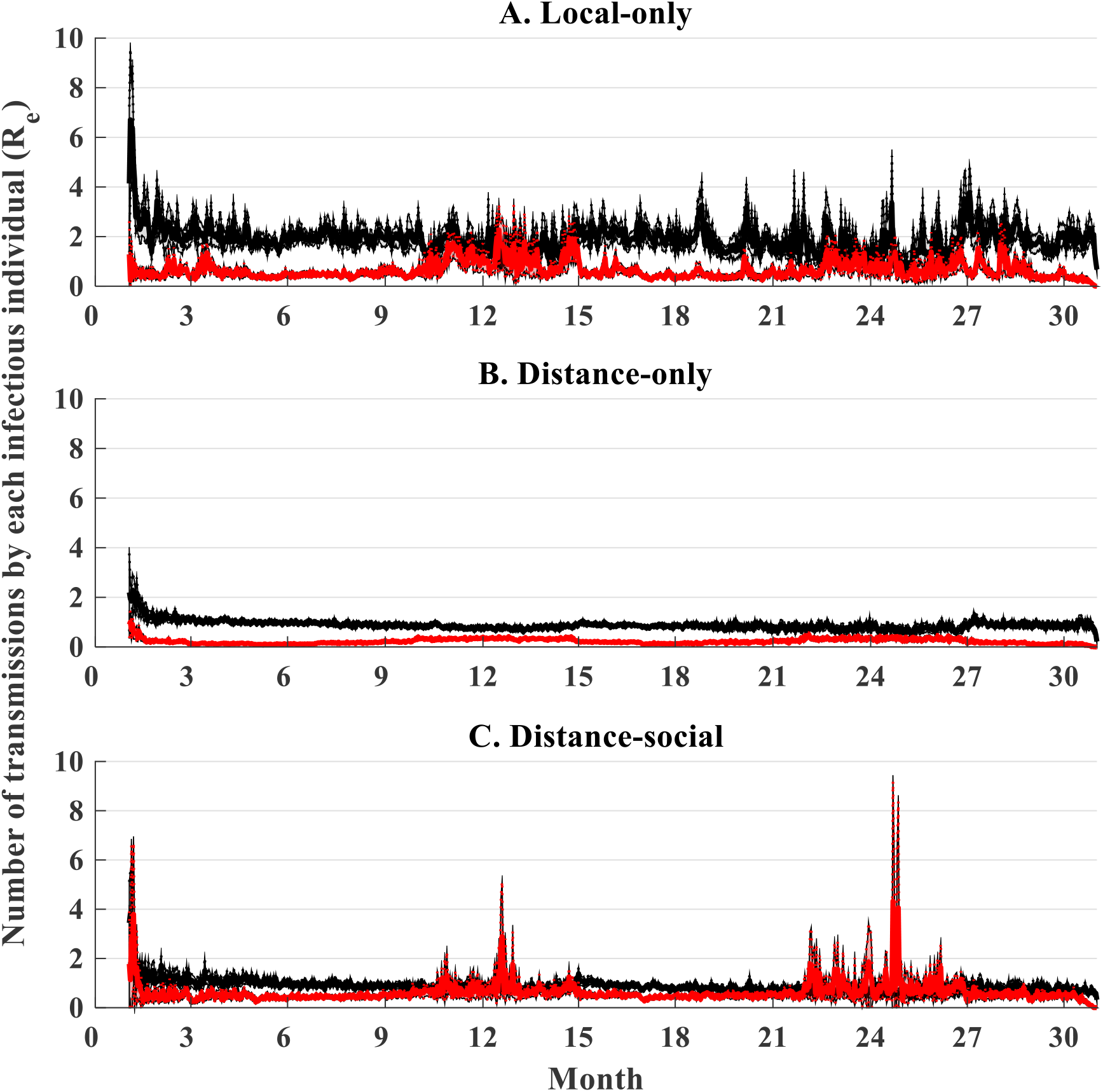
Effective reproduction number (R_e_) over time for each model. Effective reproduction number at a given time point was calculated as the average number of transmissions made throughout the infectious period for individuals that initially became infectious on day *t* (where *t* is a day on the X-axis). Dark lines are the means of 100 simulations using random samples from the posterior distributions of the fitted models; shading indicates 95% prediction intervals of the means. Overall means with 95% prediction intervals for each model and each transmission mechanism (black: direct, red: carcass-based) were: A) 1.9 [1.4, 2.3], 0.6 [0.4, 0.9], B) 0.9 [0.8 1.0], 0.2 [0.2, 0.3], C) 0.9 [0.7, 1.1], 0.6 [0.4, 0.9]. Month 1 on the x-axis is January, 2014.

## DISCUSSION

Prediction of spatial pathogen dynamics is often challenged by partially observed data, multiple pathogen introductions, and a limited understanding of host contact processes. Our approach accounted for partially observed surveillance data and re-introductions while examining how different dimensions of host contact heterogeneity (social and spatial processes) affect inference of spatial transmission. Not accounting for social structure in spatial transmission led to very different inferences of STKs and R_e_, as well as the role of sex, birth pulses, and transmission mechanisms in driving spatial spread. As STKs are used to plan resource allocation to surveillance and control, reducing uncertainty in STK estimates is crucial for maximizing efficiency and effectiveness of response plans [54]. Our results suggest that robust estimates of STKs (and other key epidemiological quantities such as R_e_) may require consideration of both social and spatial processes for accurate prediction of spatial spread. Secondly, the striking differences among model predictions in the role of carcass-based transmission and magnitude and variation in R_e_ suggests that models that exclude social structure may provide different ecological insights about which factors are most important in transmission and persistence of ASFv (e.g., carcass versus direct transmission processes, males versus females, or family groups versus independent males).

Spatially-explicit disease models often predict that local variations in group size can be extremely important in driving pathogen transmission due to threshold effects [55], but that the magnitude of these effects is context-specific. For example, an analysis that evaluated the impact of social-network organization on disease spread in 47 species suggests that outbreaks of highly contagious pathogens may be larger and persist longer in highly fragmented, gregarious species when compared to species with hierarchical social systems that are more socially connected, which in contrast, are more prone to epidemic outbreaks of low to moderately transmissible pathogens due to high connectivity [1]. This suggests that highly fragmented social organization might slow the rate of spatial disease spread. Our work supports this idea because the distance-only model predicted more rapid spatial spread relative to the distance-social model, despite similar natal dispersal distances. Also, in order for the distance-only model to fit the case data as well as possible, a larger median decay rate parameter on the distance function was estimated relative to the distance-social model, explaining the faster rate of spatial spread and less locally dense STK relative to the distance-social model. However, similar types of social behavior can impact disease spread differently [56]. For example, territorial behaviors have been shown to reduce the spread of bovine tuberculosis in European badgers to adjacent social groups [57, 58], increase the risk of macroparasite infection in African antelope [11], and play only a minor role in the spread of pathogens like feline calcivirus or canine distemper virus in Serengeti lions [56]. This suggests that social structure may not always be important for accurately capturing spatial spread dynamics, and that it is important to identify the most important dimensions (if any) of social behavior that might modify spatial spread.

Evaluating the impact of social structure on disease spread is facilitated when social interactions among hosts are known [24, 59], but in many cases social structure must be captured through representing processes such as connectivity structure or group size because individual-level interactions are unknown [60]. Previous work on raccoon rabies demonstrates the value of integrating social structure and space use (natal dispersal and home range size) into predictions of spatial disease spread and evaluating the effectiveness of control [7, 23]. Similarly, a simulation study of pathogens that are epidemiologically similar to foot-and-mouth disease and classical swine fever showed how realistic social structure can slow the spatial spread of disease and lower persistence, and that this effect is disproportionately strong for acute infections [27]. Spatial disease spread can be represented using a number of modeling frameworks (spatially-explicit deterministic models, correlation models, individual-based lattice models, and reaction-diffusion models, etc.), but methods to account for social structure and space use currently in estimates of STKs (as demonstrated here) remain limited, especially in wildlife disease systems [3, 55, 61].

One way to account for social structure and space use concurrently is to use spatial network models [17], but this approach makes it difficult to disentangle the relative effects of social versus spatial processes on pathogen transmission, which is important for optimizing control strategies (but see [60] for a novel approach that addresses this issue). In contrast, the field of movement ecology has developed new strategies to account for heterogeneities due to animal movement [3, 4] and social behavior [62, 63], but these methods remain underdeveloped for application in disease ecology [3], and present an opportunity for future work. Development of conventional methods that allow separate inference and understanding of the role of spatial and social drivers of pathogen transmission dynamics is important not only for an improved mechanistic understanding of spatial pathogen dynamics but also for improved risk assessment and optimal control strategies by highlighting which processes (e.g., host clustering, movement patterns) at what scale should be targeted for control [61].

Although IBMs can be computationally intensive, they can capture the heterogeneous traits that make each population, disease, and sampling design unique [25]. As such, they are advantageous for informing policy in many fields of epidemiology and public health that are influenced by individuals, their behaviors, and the landscape [64]. The IBM framework presented here can be used to improve predictions of spatial pathogen dynamics and risk assessment through incorporating important real-world complexities such as social clustering, sampling design, seasonal demographics, and pathogen re-introductions (see Figure 1). Though we focus on the ASFv system, this framework can be readily applied to predict spatial pathogen dynamics and understand the importance of social structure in other host-pathogen systems by using system-specific data streams. In addition to computational time, a second challenge with fitting IBMs to data can be their level of complexity and amount of stochastic variation. This can lead to wide posterior distributions on parameters, without an extremely high number of iterations (and in some cases even with a high number of iterations). As with any complex Bayesian model, this risk can be reduced through using a combination of informative prior distributions and screening results that are biologically unrealistic. During preliminary analyses we allowed more vague prior distributions on the distance decay parameter (*λ*) and found that the model could also fit the data well with a large value of *λ* but that these *λ* values were unrealistically high for wild boar movement and also led to unrealistic values for transmission probability and detection probabilities (based on expert opinion) thus we excluded these ranges in the prior distributions of our final model fitting.

The distance-social model performed best at capturing both the case and spatial spread dynamics while the distance-only model performed the worst because most spatial spread was extremely local and the distance-only model was constrained to a monotonic distribution of transmission distances. Thus, not accounting for social structure (i.e., allowing for more transmission within than between groups) while allowing for some amount of long-distance transmission (distance-only model) does a poorer job at inferring the STK because there was not enough flexibility to capture the bi- (or multi-) modal STK that best describes the spatial pathogen dynamics. Interestingly, the distance-social model performed the best and predicted the highest proportion of within-group transmission events (∼80%) while the local-only model performed the next best but predicted the lowest proportion of within-group transmission events (∼30%). This suggests that local-only model was better than the distance-only model because it restricted long-distance transmission. STKs from all models had two peaks symbolic of within-group and between-group transmission even though the local-only and distance-only models did not allow for different contact probabilities for within-versus among-group transmission because all models included spatial clustering due to family groups. Although the local-only model underestimated within-group transmission, and allowed much more between-group transmission, all of the between-group transmission was very local which allowed the model predictions to be closer to the distance-social model relative to the distance-only model. Together our results suggest that long-distance transmission is important for capturing spatial spread dynamics but that it is very rare.

Almost all transmission events for all models were within 1.5 km, with some rare events at longer distances. These STK estimates can be used to establish control and surveillance zones for wild boar in Poland. For example, carcass removal could be intensified within 1.5 km of a case detection where most transmission is occurring, with depopulation and surveillance intensified out to further distances (i.e., the tail of the STK) to contain further spread. A useful approach could be to employ an adaptive radius that focuses intervention efforts within 99% (or more – this should be validated with modeling) of the STK, but adapts surveillance based on real-time surveillance. However, the precise recommendations will depend on how soon a detection is made relative to where the infection front is currently. With ASFv travelling at 1-2 km per month [29], the radii for high-intensity culling and surveillance would need to be increased by 1-2 km for each month that detection has lagged behind the infection front, highlighting the importance of accurate predictions of spatial spread. A longer lag time for detection will also amplify challenges that arise from long-distance jumps highlighting that this process is especially important to understand. One approach to account for detection lags and anticipated spread would be to pre-determine multiple fixed radii from the STKs to delineate surveillance and control zones. This would allow for more rapid re-distribution of control resources and better targeting of surveillance resources for optimizing adaptive change based on current surveillance data.

Hunting, culling, and other anthropogenic factors could influence wild boar movement and disrupt social structure [65], thus altering STKs. While our model assumed that daily movement dynamics (contact) remained constant throughout the study period, it did account for social disruption. In cases where hunting left family group members alone, these individuals dispersed to join the nearest group (i.e., implicitly accounting for the effects of social disruption on disease spread). In terms of the potential effects of hunting or culling on daily movement, the behavioral response of wild boar to hunting activities in Europe remains nuanced, with space-use changes varying by ecological and hunting context [65]. However, feral swine culling practices in the US have shown short-term increases in movement and home range shifts, which could transiently increase ASFv transmission [66, 67], and suggests that their effects could merit further exploration.

Long-distance jumps (> 100 km from the epizootic region) were observed on multiple occasions in Poland after 2015 [37] and are thought to be due to human-mediated activities as these distances are well beyond those travelled by wild boar naturally [68, 69]. However, our model only included processes describing natural movements of wild boar (which explained most of the transmission events during our study period) because we did not have data to describe sources of human-induced long-distance transmission. Thus, developing estimates of STKs that account for mechanisms or risk factors of long-distance dispersal remains an important objective that can help target disease control efforts. Our approach allowed for some longer-distance events (on the scale of natural wild boar movements) but we assumed a monotonic functional form for contact distances, such that we did not infer the effects of spatial contact processes occurring on multiple spatial scales (beyond within-versus between-group transmission differences in the distance-social model). In order to infer spatial spread with later surveillance data (i.e., 2016-present when longer-distance events occurred multiple times), it will be important to incorporate other spatial mechanisms in the inference of the STKs, perhaps using covariate data that can inform these long-distance processes. Such an approach would provide refined recommendations for surveillance and control targets at longer distances.

We assumed that ASFv is lethal in 100% of cases, but recent evidence indicates that wild boar may survive infection [36, 81, 82]. If some hosts survive ASFv they could facilitate ASFv persistence by producing new susceptible hosts. However, the fraction of survivors is extremely small for virulent strains of ASFv (as in our study area) thus it is unclear if these individuals could realistically affect persistence. Modeling research to understand the potential role of survivors in the enzootic dynamics of ASFv could be useful for identifying conditions where survivors may alter the enzootic dynamics. We also assumed that all infections were acute (with an average infectious period of 5 days). If some individuals were to become persistently infected, those individuals would have increased opportunity to transmit the virus within and among social groups and could presumably increase the average distance of pathogen transmission because there would be more opportunity for contact among wild boar at further distances [83, 84]. However, for the virulent ASFv strains circulating in Poland, there is no evidence of persistently infected wild boar that can shed ASFv in high enough concentrations to infect other wild boar [83]. Nonetheless, surveillance for changes in virulence are important because they could change STKs over time.

Our models inferred substantial variation in R_e_ over time due to seasonally varying carcass persistence, with the distance-social model predicting the largest peaks in R_e_ due to increases in carcass and ASFv persistence in cold weather [45, 74, 75]. Prior research the Baltic States and Poland also indicated that the seasonality of ASFv cases in wild boar peaked in winter and summer [76]. Although this trend was observed across the entire European Union, these studies could not prove causality [77]. Results of our analysis suggest that carcass surveillance and removal may be especially important during the colder months due to the potentially increased role of carcass-based transmission.

Our estimates of R_e_ (ranging from 1.1-2.5 on average across models) due to different STK assumptions were similar to an estimate of R_0_ of ASFv in wild boar (1.13-3.77) in Russia [70]. However, differences between our models in estimates of R_e_ over time suggest carcass-based transmission has different roles in driving ASFv transmission, which could influence identifying the optimal control strategy considering that carcass search and removal is intensive [71]. The local-only and distance-only models suggest that carcass-based transmission is lower than direct transmission while the distance-social model suggests that the two types of transmission occurred at similar frequencies. Although carcasses have longer infectious periods than the alive phase during the cooler months, transmission probability given contact is lower and contact rates are lower relative to live individuals because carcasses do not move [45, 72, 73]. These limitations balance the advantage of potentially longer infectious periods. The distance-social models lead to higher rates of carcass-based transmission because they predict that most transmission events are extremely local. The fact that models with commonly used spatial contact structures produce different inferences of the importance of different transmission mechanisms when social structure is added emphasizes: 1) the importance of further developing inference methods that account for social structure in spatial spreading, and 2) the need for approaches that aim to reduce uncertainty in estimates of STKs, e.g., [9].

Interestingly, our models also predicted differences in the role of sex in transmission over time. The distance-social model predicted a slight female bias because this model predicted that most transmission events occurred within family groups, which are female-biased. In contrast, the local-only and distance-only models predicted a slight male bias because they predicted more between-group transmission and with a 50:50 sex ratio there are more independent males relative to family groups. Although it is known that males will travel longer distances than females [78], especially during mating season to seek out females, we did not account for this temporal heterogeneity in dispersal. Considering these types of movement heterogeneities in future work could be important for improving our understanding of which sex might present a higher risk of ASFv transmission and persistence.

Although our STK estimates are based on wild boar ecology in eastern Poland they can provide baseline guidance for planning control in other settings. Response plans in areas that are ASFv-free often rely only on local host movement ecology because there are no local data on how ASFv might transmit. Our estimates of STKs suggest that transmission of ASFv in wild boar (or pigs) in other ecological settings might mainly occur over the closer daily movement estimates within home ranges due to the social processes in this host species. To further refine STK estimates for specific ecological contexts, parameters for the local demographic (e.g. litter size, age at reproductive maturity, gestation time, longevity, seasonal birth patterns, landscape carrying capacity), movement (e.g. natal dispersal distance and timing, daily home range exploration), and social processes (maximum group size) can be used as inputs in our simulation model with the Poland-derived estimates for transmission rates to explore how local ecological factors might affect STKs or their uncertainty.

Considered separately, spatial and social processes can have similar impacts on pathogen transmission dynamics. For example, social aggregation (e.g., family groups, herd living) and spatial structuring can both restrict the spread of pathogens [1, 79]. However, our results highlight that spatial and social processes can also have quite different impacts on epidemiological quantities, especially estimates of STKs, R_e_, the frequency of different transmission mechanisms, and potential risk factors such as sex. Being able to appropriately infer the role of these quantities is crucial for optimizing disease control strategies. When these contact heterogeneities are inappropriately accounted for it can bias inference (e.g., [80]) and potentially misguide policy decisions [54]. Moving forward, the field of disease ecology should emphasize the development and use of methods that account for spatial and social contact processes separately so that their relative roles in driving pathogen transmission dynamics can be inferred and understood. This will allow uncertainties in contact processes to be appropriately evaluated and incorporated into predictions of spatial spread [54], for monitoring designs to be optimized, and for risk factors to be identified more accurately so that controls can be targeted to the most important risk factors.

## Acknowledgements

We would like to thank 5 anonymous reviewers that provided helpful comments that improved clarity of the manuscript. KMP was supported by the United States Department of Agriculture, Animal and Plant Health Inspection Service’s National Feral Swine Damage Management Program. AG was supported by the United States Department of Agriculture, Animal and Plant Health Inspection Service’s APHIS Science Fellowship. TP was supported by the National Science Centre, Poland (grant number 2014/15/B/NZ9/01933) and Ministry of Agriculture, Czech Republic (grant number QK1910462). We thank M. Łyjak, A. Kowalczyk, K. Śmietanka, and G. Woźniakowski from Department of Swine Diseases, National Veterinary Research Institute in Pulawy, Poland, for surveillance data. N. Selva provided valuable information on carcass persistence time.

## Supplemental Material

### Contents

**Table S1.** Model selection, goodness of fit, and prior and posterior distributions.

| Table S1. | Contact structure: | Local-only | Distance-only | Distance-social |
| --- | --- | --- | --- | --- |
| | $R^2$ | | | |
| Monthly cases<br>(Median of the $R^2$'s for 1000 individual time series ± 95% confidence interval) | in sample<br>all | $0.57 \pm 0.0059$<br>$0.48 \pm 0.0054$ | $0.51 \pm 0.0054$<br>$0.43 \pm 0.0046$ | $0.55 \pm 0.0060$<br>$0.47 \pm 0.0049$ |
| Monthly distance from border<br>(Median of the $R^2$'s for 1000 individual time series ± 95% confidence interval) | in sample<br>all | $0.28 \pm 0.018$<br>$0.19 \pm 0.016$ | $0.29 \pm 0.017$<br>$0.14 \pm 0.011$ | $0.31 \pm 0.018$<br>$0.28 \pm 0.011$ |
| Monthly cases<br>($R^2$ of median values from 1000 points at each month) | in sample<br>all | 0.65<br>0.55 | 0.57<br>0.48 | 0.63<br>0.53 |
| Monthly distance from border<br>($R^2$ of median values from 1000 points at each month) | in sample<br>all | 0.53<br>0.49 | 0.49<br>0.38 | 0.53<br>0.55 |
|  | Distance metrics<sup>a</sup> |  |  |  |
| Median absolute error in monthly live cases ± 95% confidence interval | in sample | $20 \pm 0.8$ | $20 \pm 0.9$ | $20 \pm 0.5$ |
| Median absolute error in monthly carcass cases ± 95% confidence interval | in sample | $44 \pm 1.4$ | $43 \pm 2.8$ | $44 \pm 0.8$ |
| Median absolute error in monthly distance from border ± 95% confidence interval | in sample | $109.5 \pm 5.0$ | $111.7 \pm 5.7$ | $106.6 \pm 3.0$ |
| Number of values in posterior distribution | | $16 / 1,959,184 = 0.00082\%$ | $7 / 1,959,184 = 0.00036\%$ | $53 / 1,959,184 = 0.0027\%$ |
|  | Uniform Priors | 95% credible intervals of posterior distributions |  |  |
| Probability of direct transmission given proximity ($\beta^d$) | [0.0001,1]<sup>b</sup> | [0.0046, 1.0] | [0.039, 0.15] | [0.0003, 0.18] |
| Probability of carcass-based transmission given proximity ($\beta^c$) | [0.0001,1]<sup>b</sup> | [0.0002, 0.53] | [0.0038, 0.029] | [0.0001, 0.029] |
| Annual frequency of spillover from neighboring country ($\phi$) | [0,60]<sup>c</sup> | [13, 59] | [9, 60] | [6, 59] |
| Detection probability in hunted boar samples ($\rho^h$) | [0.0005,0.1]<sup>b</sup> | [0.0006, 0.012] | [0.0007, 0.021] | [0.0007, 0.017] |
| Detection probability in carcass samples ($\rho^c$) | [0.0005,0.8]<sup>b</sup> | [0.012, 0.080] | [0.017, 0.045] | [0.014, 0.30] |
| Scaling parameter on seasonal trends in carcass persistence ($\pi$) | [0.1,1.5] | [0.58, 1.47] | [0.77, 1.41] | [0.17, 1.47] |
| Scaling parameter on seasonal trends in conception probability ($\theta$) | [0.5,6] | [0.93, 5.73] | [1.31, 5.87] | [0.67, 5.88] |
| Constant contact radius ($\zeta$) | [0.5,5] | [1.09, 3.35] | NA | NA |
| Rate parameter for decay of contact probability with distance ($\lambda$) | [0.1,2.5] | NA | [1.62, 2.49] | [0.05, 2.49] |
| Probability of direct transmission given contact is in the same group ($\beta^{w,d}$) | [0.01,1] | NA | NA | [0.077, 0.98] |
| Probability of carcass-based transmission given contact is in the same group ($\beta^{w,c}$) | [0.001,1] | NA | NA | [0.14, 0.97] |

<sup>a</sup>Median distance metrics ± 95% confidence intervals for 1000 simulations from the posterior distribution.
<sup>b</sup>These prior distributions were sampled on a natural log scale.
<sup>c</sup>0 indicates one introduction ever whereas values $> 0$ indicate the number of introductions / year.

**Figure S1.**
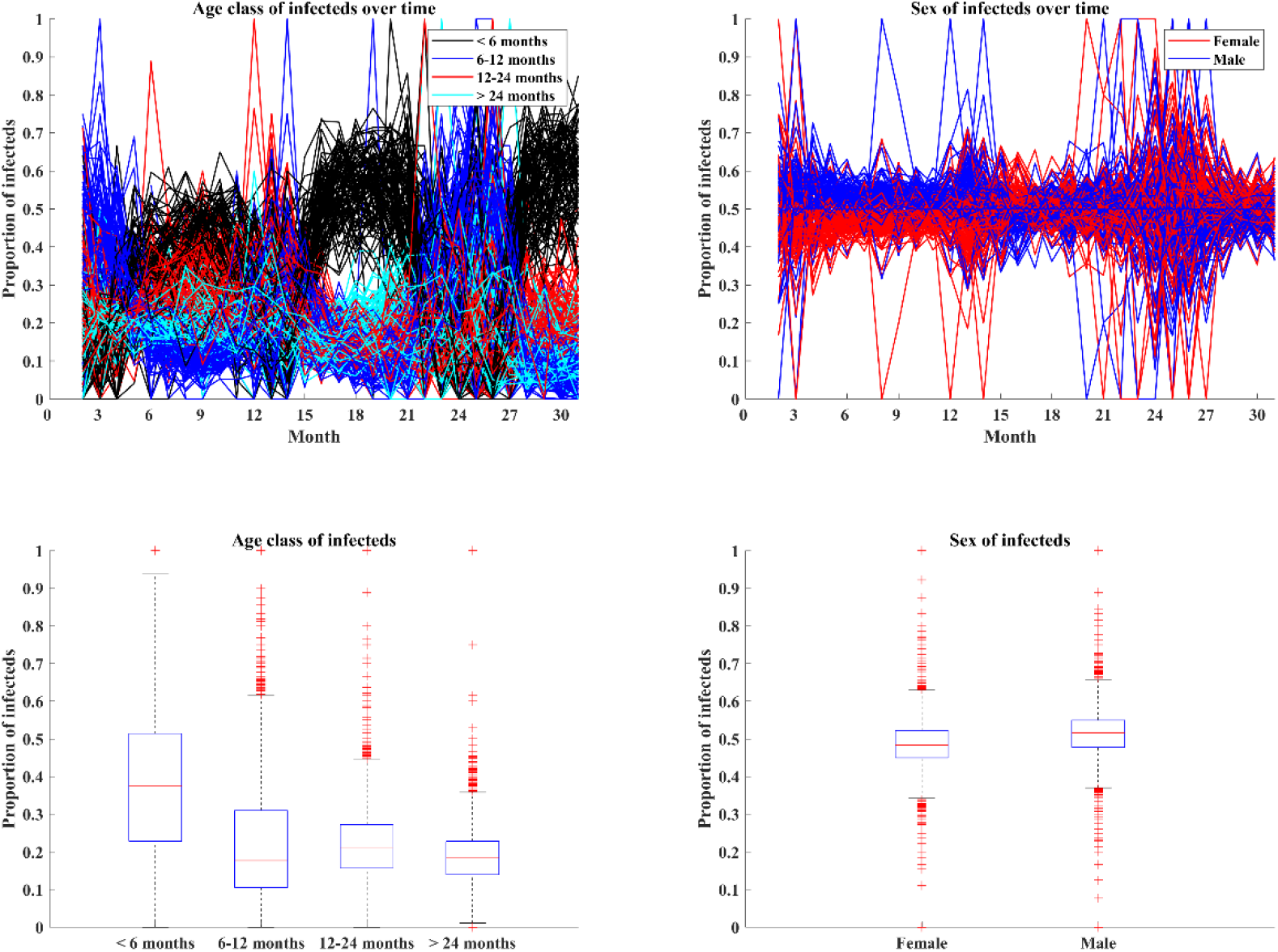
The demographic dynamics of African swine fever virus based on the local-only model. Top plots show the frequency of infection in wild boar of different age classes (left) and sexes (right). Each line is a prediction from a separate sample of the posterior distribution of the fitted exponential decay & social structure model (100 trajectories in total). Bottom plots show the corresponding age class and sex distributions of infection over all time.

**Figure S2.**
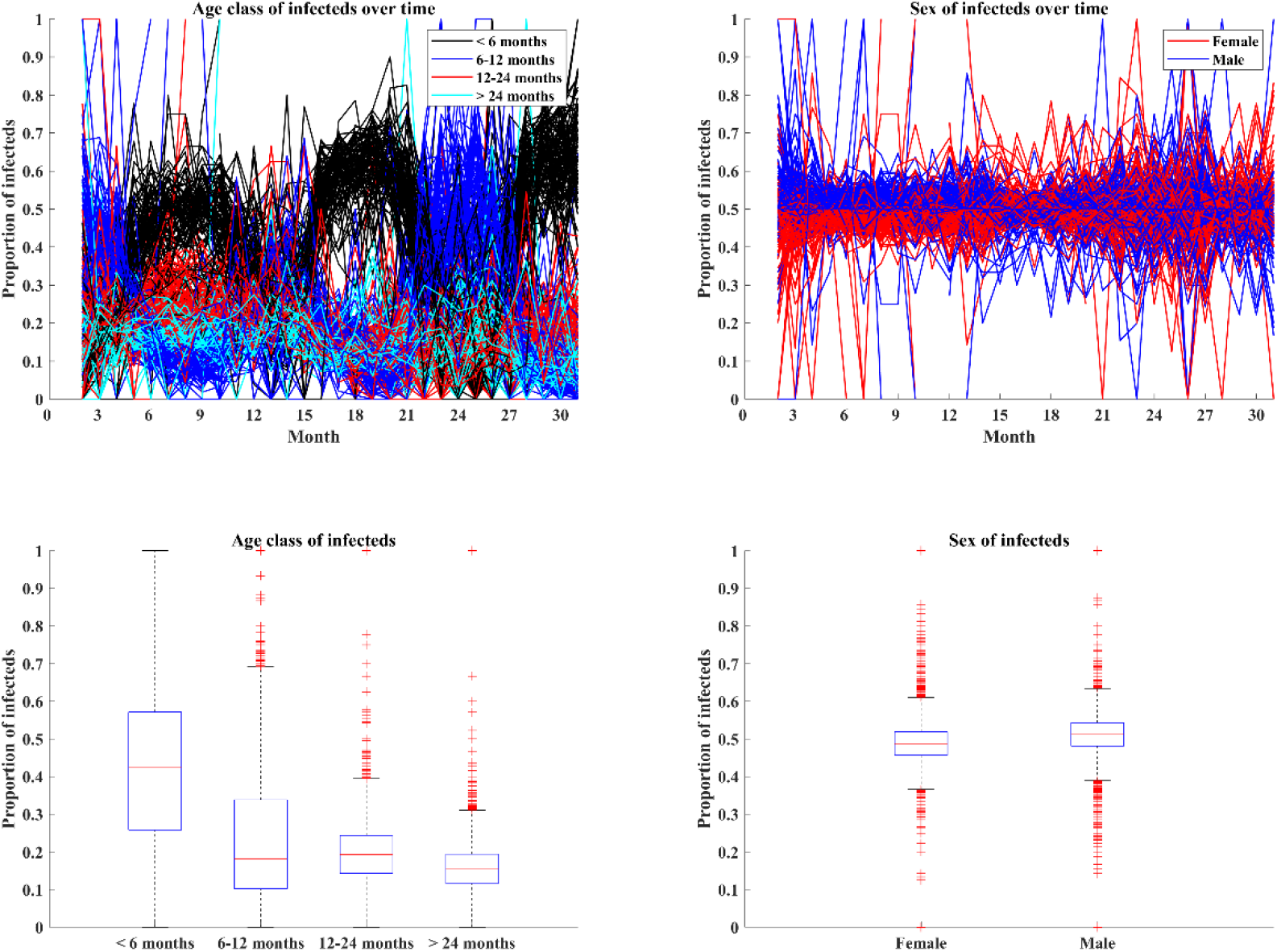
The demographic dynamics of African swine fever virus based on the distance-only model. Top plots show the frequency of infection in wild boar of different age classes (left) and sexes (right). Each line is a prediction from a separate sample of the posterior distribution of the fitted exponential decay & social structure model (100 trajectories in total). Bottom plots show the corresponding age class and sex distributions of infection over all time.

**Figure S3.**
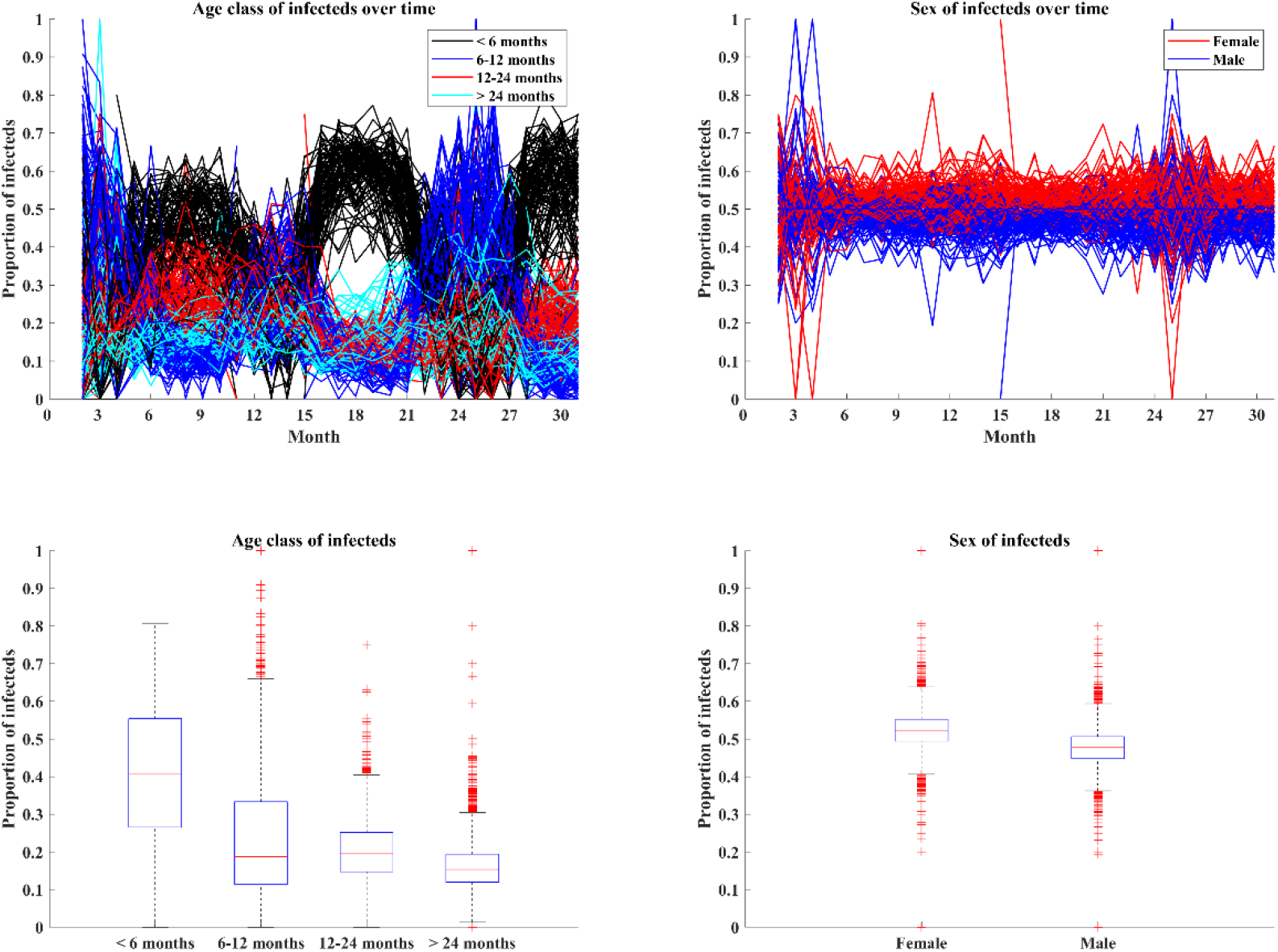
The demographic dynamics of African swine fever virus based on the distance-social model. Top plots show the frequency of infection in wild boar of different age classes (left) and sexes (right). Each line is a prediction from a separate sample of the posterior distribution of the fitted exponential decay & social structure model (100 trajectories in total). Bottom plots show the corresponding age class and sex distributions of infection over all time.

